# Bisphenol S and bisphenol F are less disruptive to cardiac electrophysiology and potentially safer for use in medical products, as compared to bisphenol A

**DOI:** 10.1101/2021.02.16.431039

**Authors:** Tomas Prudencio, Luther Swift, Devon Guerrelli, Blake Cooper, Marissa Reilly, Nina Ciccarrelli, Jiansong Sheng, Rafael Jaimes, Nikki Gillum Posnack

**Author notes:** **Corresponding author**: Nikki Gillum Posnack, Ph.D., Sheikh Zayed Institute, 6^th^ floor, M7707, 111 Michigan Avenue, NW, Washington, DC, USA 20010.

## Abstract

**Background:** Bisphenol A (BPA) is a high-production volume chemical that is commonly used to manufacture consumer and medical-grade plastic products. Due to its ubiquity, the general population can incur daily environmental exposure to BPA, while heightened BPA exposure has been reported in intensive care patients and industrial workers. Due to health concerns, structural analogues are being explored as replacements for BPA.

**Objective:** This study aimed to examine the direct nongenomic effects of BPA on cardiac electrophysiology and compare its safety profile to recently developed alternatives, including BPS (bisphenol S) and BPF (bisphenol F).

**Methods:** Whole-cell voltage-clamp recordings were performed on cell lines transfected with Nav1.5, hERG, or Cav1.2. Results of single channel experiments were validated by conducting electrophysiology studies on human induced pluripotent stem cell-derived cardiomyocytes (hiPSC-CM) and intact, whole heart preparations.

**Results:** Of the chemicals tested, BPA was the most potent inhibitor of both fast (I_Na-P_) and late (I_Na-L_) sodium channel (IC_50_ = 55.3 and 23.6 μM, respectively), L-type calcium channel (IC_50_ = 30.8 μM) and hERG channel current (IC_50_ = 127 μM). The inhibitory effects of BPA and BPF on L-type calcium channels were supported by microelectrode array recordings, which revealed shortening of the extracellular field potential (akin to QT interval). Further, BPA and BPF exposure impaired atrioventricular conduction in intact, whole heart experiments. BPS did not alter any of the cardiac electrophysiology parameters tested.

**Discussion:** Results of this study demonstrate that BPA and BPF exert an immediate inhibitory effect on cardiac ion channels, and that BPS may be a safer alternative. Intracellular signaling or genomic effects of bisphenol analogues were not investigated; therefore, additional mechanistic studies are necessary to fully elucidate the safety profile of bisphenol analogues on the heart.

## INTRODUCTION

Bisphenol A (BPA) is a high production volume chemical, with roughly 8 million metric tons used each year to manufacture polycarbonate plastics (e.g., food and beverage containers, medical devices), epoxy resins (e.g., aluminum can liners), and in thermal printing applications (PR newswire 2016; Shelby 2008). Human exposure to BPA can occur daily, and as a result, biomonitoring studies have detected BPA in 91-99% of the general population (Calafat et al. 2005; Chen et al. 2016a; Lehmler et al. 2018; Vandenberg et al. 2007, 2010). Although environmental exposure to BPA occurs at a relatively low dose (Koch and Calafat 2009; Vandenberg et al. 2007, 2010), occupational (Hines et al. 2018; Ribeiro et al. 2017) and clinical environments can result in exceedingly high BPA exposure (Calafat et al. 2009; Duty et al. 2013; Gaynor et al. 2018; Huygh et al. 2015; Testai et al. 2016). Indeed, BPA was detected in 60% of neonatal intensive care unit (NICU) supplies, including items used for feeding, bandages, breathing support, intravenous and parenteral infusion (Iribarne-Durán et al. 2019). Clinical exposure can also result in heightened and/or prolonged BPA exposure in young patients, due to an underdeveloped metabolic system (Calafat et al. 2009). In the NICU setting, premature infants had urinary BPA levels that ranged from 1.6–946 μg/L (Calafat et al. 2009) and the degree of exposure was linked to high-intensity treatment that required multiple (plastic) medical devices (Duty et al. 2013). Similarly, adult ICU patients were found to have urinary BPA levels that ranged from 6.1–680 μg/L when undergoing extracorporeal membrane oxygenation in conjunction with continuous vevo-venous hemofiltration (Huygh et al. 2015).

BPA exposure is concerning, particularly in sensitive patient populations, as accumulating evidence suggests that BPA exerts a negative impact on cardiovascular health (Bae et al. 2012; Bae and Hong 2015; Han and Hong 2016; Melzer et al. 2010, 2012). A 10-year longitudinal study found that BPA exposure was associated with a 46-49% higher hazard ratio for cardiovascular and all-cause mortality (Bao et al. 2020). Further, epidemiological studies have reported associations between BPA exposure and an increased risk of myocardial infarction, hypertension, coronary and peripheral artery disease, and a decrease in heart rate variability (reviewed previously (Posnack 2014; Ramadan et al. 2020). Experimental studies have noted that BPA exposure can antagonize ion channels, impair electrical conduction, and precipitate triggered arrhythmias (Belcher et al. 2011; Deutschmann et al. 2013; Feiteiro et al. 2018; Michaela et al. 2014; Posnack et al. 2015; Wang et al. 2011; Yan et al. 2011). *In vitro* studies performed in HEK, neuronal, and smooth muscle cells have shown that BPA inhibits T-type and L-type calcium channel current (Deutschmann et al. 2013; Feiteiro et al. 2018; Michaela et al. 2014). In cardiac tissue, such an alteration in calcium channel current would alter nodal cell depolarization, atrioventricular conduction, and the plateau phase of the cardiac action potential. Further, BPA exposure can disrupt intracellular calcium handling, resulting in calcium leak from the sarcoplasmic reticulum and an increased propensity for triggered arrhythmias (Gao et al. 2013; Liang et al. 2014; Ramadan et al. 2018). Of interest, BPA exposure was observed to increase calcium-mediated triggered activity and ventricular arrhythmias in females (but not males) that were subjected to catecholamine stress. Notably, such alterations in calcium handling were attenuated in an estrogen-receptor knockout model (Yan et al. 2011), which supports the claim that BPA-induced effects are sex specific.

With increasing health concerns, structurally similar chemicals are being explored as replacements for BPA (Chen et al. 2016a). Two such substitutes, bisphenol S (BPS) and bisphenol F (BPF), are used to manufacture consumer products that don ‘BPA-free’ labeling. For example, BPS is used to produce polyethersulfone plastic food containers, medical-grade products, epoxy resins, and is found in thermal printing applications (Chen et al. 2016a; Lehmler et al. 2018). Unfortunately, many of these alternative chemicals are considered ‘regrettable substitutions’, as BPS and BPF may exert biological effects that are similar to BPA (Kojima et al. 2019; Moon 2019; Trasande 2017). To date, it is unclear whether BPA alternatives offer a superior cardiac safety profile, as recent work suggests that BPS and BPF may also impair cardiac function (Ferguson et al. 2019; Gao et al. 2015; Mu et al. 2019). Recent biomonitoring studies have observed an uptick in BPS and BPF exposure in the general population as BPA is phased out and replaced (Lehmler et al. 2018), which highlights the urgent need to investigate the impact of BPA analogues on cardiac health.

We compared the cardiac safety profile of BPA, BPS, and BPF using whole-cell voltage clamp experiments to identify the half maximal inhibitory concentration (IC_50_) of four key cardiac ion channels, highlighted by the CiPA (comprehensive in vitro proarrhythmia) initiative (Colatsky et al. 2016; Sager et al. 2014). The results of single channel experiments were validated by conducting electrophysiology studies on human induced pluripotent stem cell-derived cardiomyocytes (hiPSC-CM) using microelectrode array (MEA) recordings. Importantly, hiPSC-CM have been widely adopted as a tool for preclinical safety testing to measure alterations in cardiac automaticity, conduction velocity, depolarization, and repolarization time (Chen et al. 2016b). To aid in the translation of our *in vitro* cell studies, we employed an intact, whole heart preparation for direct assessment of cardiac electrophysiology. We hypothesized that inhibitory effects of bisphenol chemicals on calcium current would present with delayed atrioventricular conduction in cardiac preparations. Further, we hypothesized that BPS and/or BPF exposure would have less effect on cardiac electrophysiology, due to differences in chemical structure that may increase the potency of BPA for voltage-gated channels (Deutschmann et al. 2013).

## METHODS

### Reagents

Bisphenol A (CAS #80-05-7), bisphenol S (CAS #80-09-1), and bisphenol F (CAS #620-92-8) were purchased from Sigma Aldrich (≥98% purity, analytical standard). Stock solutions of BPA, BPS, or BPF were prepared in 99+% dimethyl sulfoxide (DMSO), and working concentrations were prepared directly in cell culture media (voltage-clamp recordings, MEA studies) or Krebs-Henseleit (KH) crystalloid buffer (intact heart preparations) to obtain a final concentration between 0.01-100 μM BPA, BPS, or BPF. This range of doses was selected to mimic environmental, clinical, and supraphysiological exposure levels (Ramadan et al. 2018).

### Whole-cell voltage-clamp recordings

Nav1.5, Cav1.2, and hERG channel recordings were performed at room temperature (25°C) using stably transfected cell lines, as previously described (Jaimes et al. 2019). For *<u>Nav1.5 recordings</u>*, the extracellular solution included 137 mM NaCl, 10 mM dextrose, 10 mM HEPES, 4 mM KCl, 1 mM MgCl_2_, and 1 mM CaCl_2_. The intracellular solution consisted of 120 mM CsOH, 120 mM aspartic acid, 10 mM EGTA, 10 mM CsCl, 10 mM HEPES, 5 MgATP, and 0.4 mM Tris-GTP. The voltage protocol was approximately 1 sec in duration, repeated at 0.1 Hz. Sodium channel recordings were performed using HEK293 cells transfected with Nav1.5 cDNA. Cells were repolarized from −95 to −120 mV for 200 msec, depolarized from −120 to −15 mV for 40 msec, and then further depolarized to +40 mV for 200 msec. This was followed immediately by a voltage ramp down phase from +40 to −95 mV for 100 msec. ATXII (20 nmol/L) was included in the extracellular solution to induce Nav1.5 late current, as previously described (Mantegazza et al. 1998). Tetrodotoxin (30 μM) was applied at the end of each recording to determine the current baseline. *<u>Cav1.2 recordings</u>* were performed using CHO cells stably transfected with Cav1.2 cDNA. Cells were depolarized from −80 mV to 0 mV for 40 msec, further depolarized to +30 mV for 200 msec, followed by a voltage ramp down phase from +30 mV to −80 mV for 100 msec. Recording stability was assessed by applying the voltage protocol in control solution for 12 consecutively recorded traces with <10% difference. *<u>hERG recordings</u>* were performed using HEK293 cells stably transfected with hERG cDNA. The extracellular solution included 130 Mm NaCl, 12.5 mM dextrose, 10 mM HEPES, 5 mM KCl, 1 mM MgCl_2_:6H_2_O, and 1 mM CaCl_2_. The intracellular solution consisted of 120 K-gluconate, 20 mM KCl, 10 mM HEPES, 5 EGTA, and 1.5 MgATP. The voltage protocol was 5 sec in duration, repeated at 0.1 Hz. Cells were depolarized −80 mV to +40 mV for 500 msec, followed by a voltage ramp down phase from +40 mV to −80 mV for 100 msec. A hERG potassium channel blocker (10 μM E-4031) was applied at the end of each recording to determine the baseline. Recordings were collected before and after bisphenol chemical exposure; chemical potency was calculated by dividing the steady state current amplitude by the average amplitude from the last 5 traces measured in control solution to calculate the fractional block. This was plotted against the bisphenol chemical concentration tested, fitted with the Hill Equation to generate a half-maximal inhibitory concentration (IC_50_) and the Hill coefficient.

### Human cardiomyocyte microelectrode array recordings

hiPSC-CM (iCell cardiomyocytes^2^, female donor #01434, Fujifilm) were plated onto fibronectin-coated microelectrode arrays at a density of 50-75,000 cells/well (24-well plate, Axion). Cells were defrosted in iCell cardiomyocyte plating media in a cell culture incubator (37°C, 5% CO_2_) for 2 hours, thereafter cells were cultured in iCell maintenance media for the duration of study. Treatment groups included 0.01% DMSO (vehicle), 0.01-100 μM BPA, BPS, or BPF. Cells were treated for 15 minutes, and then extracellular field potential signals were recorded in response to external stimulation (1-2 Hz). Extracellular field potential duration (FPD) was measured and rate corrected with Frederica formula (FPDc). Disturbances in the recorded waveform can be used to predict the identity of ion channels impacted by chemical exposure, with FPD analogous to an *in vitro* QT interval that correlates with action potential duration at 50% repolarization (Asakura et al.; Clements 2016).

### Animals

Animal protocols were approved by the Institutional Animal Care and Use Committee at Children’s National Research Institute and followed the National Institutes of Health’s *Guide for the Care and Use of Laboratory Animals*. Bisphenol chemicals are xenoestrogens that may cause exaggerated cardiac effects in females (Ben-Jonathan and Steinmetz 1998; Yan et al. 2011); accordingly, experiments were performed using female Sprague-Dawley rats, aged 3-4 months (Taconic Biosciences, strain NTac:SD, from NIH Genetic Resource stock, n=66). Animals were housed in conventional acrylic rat cages in the Research Animal Facility, under standard environmental conditions (12:12 hour light:dark cycle, 18–25°C, 30-70% humidity). Each animal served as its own control, with electrophysiology measurements collected at baseline and again after treatment.

### Intact heart preparations

Animals were anesthetized with 3% isoflurane; the chest was opened, the heart was rapidly excised, and the aorta was cannulated. The isolated, intact heart was then transferred to a temperature-controlled (37°C) constant-pressure (70 mmHg) Langendorff-perfusion system. Excised hearts were perfused with a modified Krebs-Henseleit buffer bubbled with carbogen, as previously described (Jaimes et al. 2019). Pseudo-electrocardiograms (ECG) were recorded in lead II configuration, and biosignals were acquired in iox2 and analyzed in ecgAUTO. Isolated hearts remained stable with minimal fluctuations in heart rate or electrophysiology parameters following 0.01% DMSO media perfusion (vehicle; **Figure 1**). To account for animal variability, ECG recordings were collected throughout the study, during control media perfusion (15 min) and in response to bisphenol chemical exposure (15 min). Similarly, electrophysiology measurements (see below) were measured at baseline, after 15 min chemical exposure, and again after 15 min washout with KH media.

**Figure 1.**
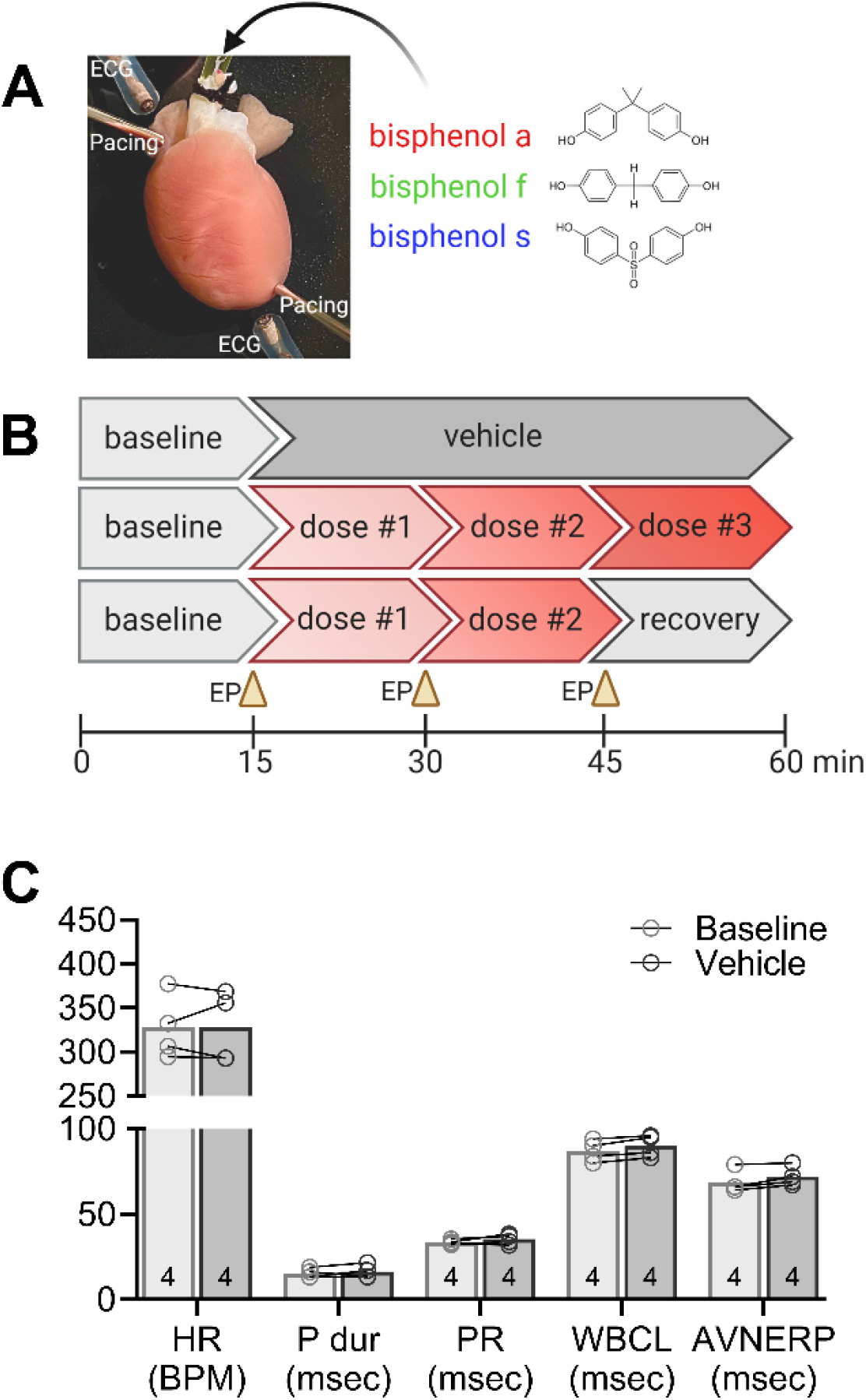
Experimental design and vehicle control parameters. **(A)** Langendorff-perfused rat heart shown with pacing electrodes on the right atria and apex, and monopolar electrodes placed to record electrocardiograms. **(B)** Schematic timeline depicts perfusion protocols used in the study, including (top) vehicle control exposure, (middle) bisphenol chemical dose response, (bottom) bisphenol chemical dose response and subsequent recovery. **(C)** Cardiac electrophysiology parameters are consistent over time, and similar between baseline and vehicle (0.01% DMSO) exposure. Values reported as mean ± SD. Statistical significance determined by RM-ANOVA with multiple comparisons testing (0.1 FDR). Number of replicates indicated in each bar graph (n=4). ECG = location of electrocardiogram electrode, EP = electrophysiology protocol, HR = heart rate, BPM = beats per minute, P dur = P wave duration, PR = PR interval, WBCL = Wenckebach cycle length, AVNERP = atrioventricular node effective refractory period, msec = milliseconds

### Electrophysiology measurements

A pacing electrode was positioned on the right atrium for assessment of atrioventricular (AV) conduction time, AV node refractory period (AVNERP) and Wenckebach cycle length (WBCL). WBCL was defined as the shortest S1-S1 pacing interval that resulted in 1:1 atrioventricular conduction. AVNERP was defined as the shortest S1-S2 pacing interval that resulted in 1:1 atrioventricular conduction. Electrophysiology studies were performed using a Bloom Classic electrophysiology stimulator (Fisher Medical) set to a pacing current 1.5x the minimum pacing threshold, with 1 msec monophasic pulse width. For each parameter, the pacing cycle length (PCL) was decremented to pinpoint the PCL before loss of capture was observed.

### Statistical Analysis

Results are reported as mean ± standard deviation. Data normality was assessed by Shapiro-Wilk testing (Graphpad Prism). Statistical analysis was performed using one-way analysis of variance (ANOVA) for microelectrode array recordings or two-way analysis of variance with repeated measures (RM-ANOVA) to compare baseline vs treatment in whole heart experiments. Significance was defined by an adjusted p-value (q<0.1) after multiple comparisons testing with a false discovery rate of 0.1; significance is denoted in the figures with an asterisk (*).

## RESULTS

### BPA exerts a greater inhibitory effect on ion channels, compared with BPS or BPF

Whole-cell voltage clamp recordings were performed on cells transfected with one of four cardiac ion channels, as highlighted by CiPA (Colatsky et al. 2016; Sager et al. 2014). Currents were evoked and recorded before and after exposure to BPA, BPS, or BPF, and a half-maximal inhibitory concentration (IC_50_) was computed by testing the respective affinities to each ion channel. Collectively, BPA had the highest affinity for each ion channel tested, compared to both BPF and BPS, and current suppression was concentration dependent (**Figure 2**,**3**). Peak sodium current (I_Na-P_) was suppressed with an IC_50_ of 55.3 μM BPA, 232 μM BPF, and 1090 μM for BPS; late sodium current (I_Na-L_) was suppressed at lower doses, with a computed IC_50_ of 23.6 μM BPA, 100 μM BPF, and 369 μM BPS (**Figure 2A**,**B**). In ventricular tissue, I_Na-P_ is responsible for action potential upstroke (phase 0) and I_Na-L_ is involved in the plateau phase (phase 2). Accordingly, inhibition of I_Na-P_ is likely to slow depolarization and electrical conduction, while inhibition of I_Na-L_ can shorten the action potential duration. L-type calcium channel current (I_CaL_) was also the most sensitive to BPA exposure, with an IC_50_ of 30.8 μM, compared to 76 μM BPF and 333 μM BPS (**Figure 3A**). In ventricular myocytes, calcium current (I_CaL_) plays a prominent role in the plateau phase, and also contributes to the action potential upstroke in nodal cells. Inhibition of I_CaL_ can slow sinus rate, delay atrioventricular conduction, and shorten the ventricular myocyte action potential. Finally, the rapid delayed rectifier potassium current (I_Kr_) was suppressed at higher bisphenol concentrations, with a measured IC_50_ of 127 μM BPA, 209 μM BPF, and 633 μM BPS (**Figure 3B**). Bisphenol exposure could suppress I_Kr_ and prolong cardiac repolarization (phase 3) at high concentrations.

**Figure 2.**
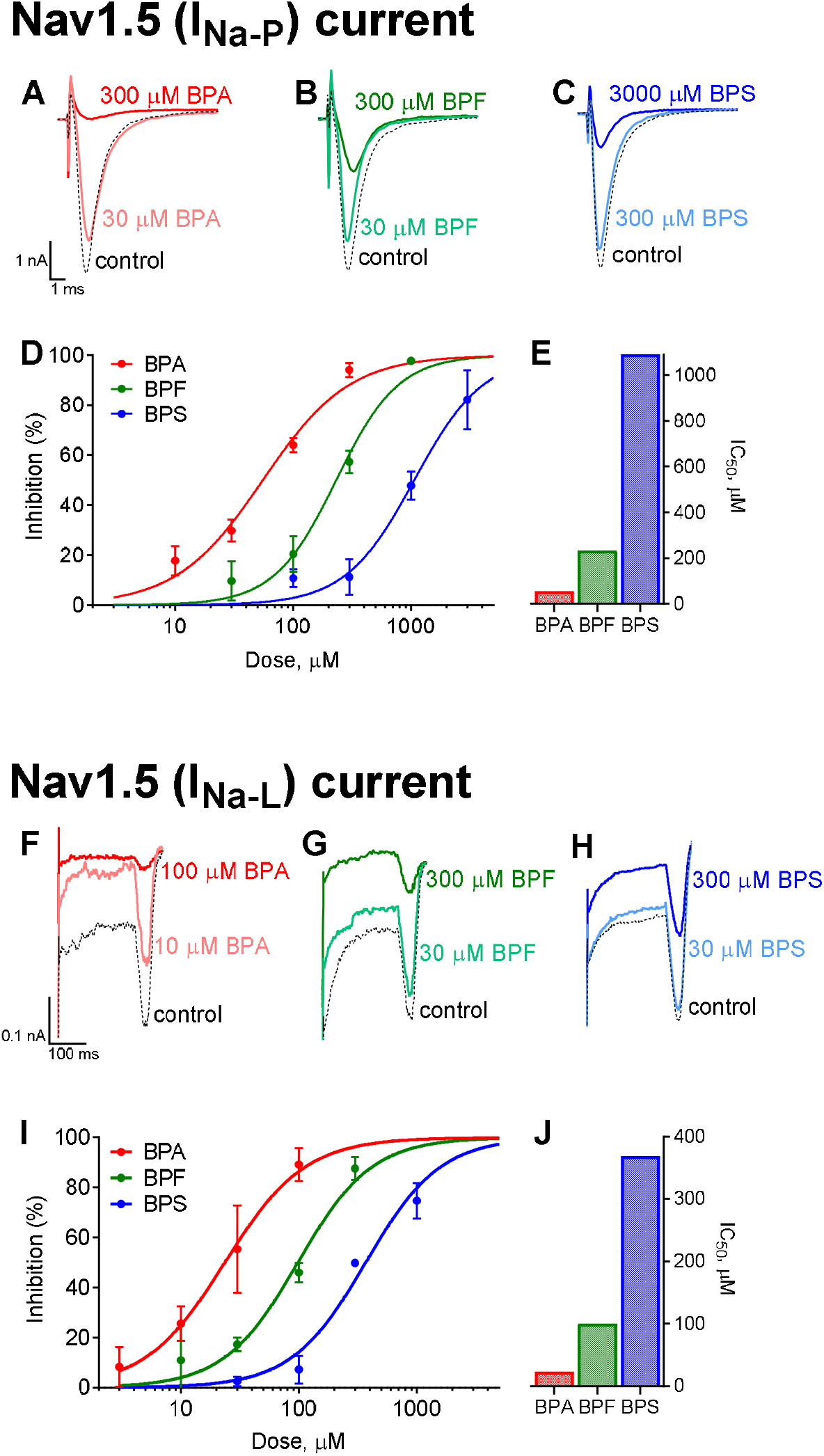
Bisphenol inhibition of sodium currents. Whole-cell voltage clamp recordings of fast/peak sodium current (I_Na-P_) following exposure to **(A)** BPA, **(B)** BPF, or **(C)** BPS. **(D)** Dose-dependent inhibition of I_Na-P_ (mean ± SD). **(E)** Calculated IC_50_ values are shown. Whole-cell voltage clamp recordings of late sodium current (I_Na-L_) following exposure to **(F)** BPA, **(G)** BPF, or **(H)** BPS. **(I)** Dose-dependent inhibition of I_Na-L_ (mean ± SD). **(J)** Calculated IC_50_ values are shown.

**Figure 3.**
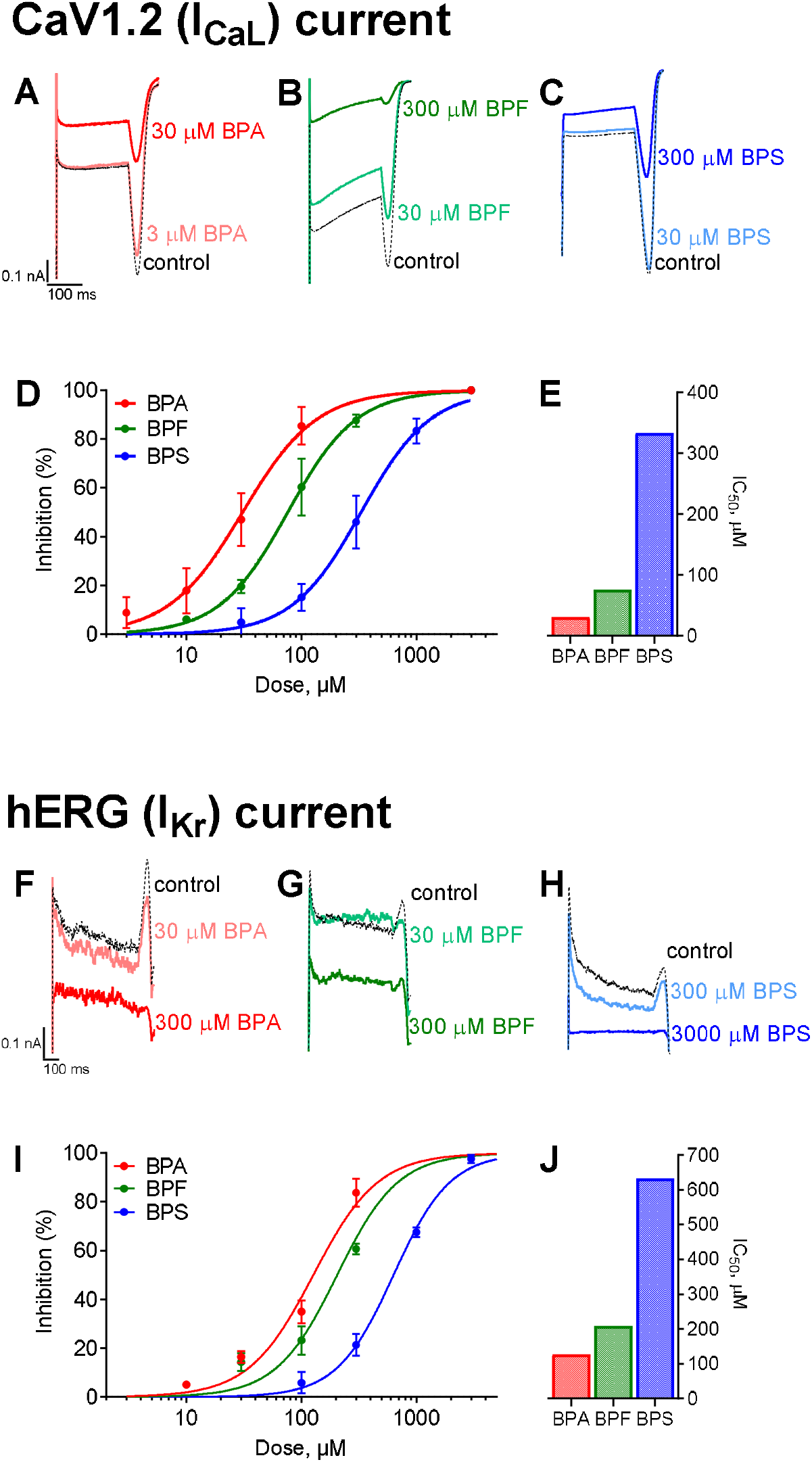
Bisphenol inhibition of calcium and potassium currents. Whole-cell voltage clamp recordings of L-type calcium current (I_CaL_) following exposure to **(A)** BPA, **(B)** BPF, or **(C)** BPS. **(D)** Dose-dependent inhibition of I_CaL_ (mean ± SD). **(E)** Calculated IC_50_ values are shown. Whole-cell voltage clamp recordings of hERG current (I_Kr_) following exposure to **(F)** BPA, **(G)** BPF, or **(H)** BPS. **(I)** Dose-dependent inhibition of I_KrL_ (mean ± SD). **(J)** Calculated IC_50_ values are shown.

### BPA and BPF exposure alters the extracellular field potential of human cardiomyocytes

To validate the effects of bisphenol chemicals on cardiac electrophysiology, we employed hiPSC-CM that express key cardiac ion channels (Edwards and Louch 2017). hiPSC-CM were cultured atop microelectrodes and extracellular field potentials were recorded (**Figure 4A, D, G**), with FPD measurements analogous to an *in vitro* QT interval that correlates with the action potential duration (Asakura et al.; Clements 2016). Acute BPA exposure resulted in a slight non-monotonic dose response (**Figure 4B**), wherein no effect on FPDc was observed at the lowest BPA dose tested (0.01 μM) and a 7.5% increase in FPDc was observed at 100 nM BPA (q<0.005). Notably, low dose effects have previously been reported for BPA and other endocrine-disrupting chemicals that can present with a non-monotonic dose response (Birnbaum 2012; Vandenberg 2014). At higher BPA doses, FPDc shortened significantly compared to the vehicle (13.8% at 30 µM, 37.3% at 100 µM, q<0.0001). Low dose effects were not observed for either BPF or BPS (**Figure 4E, H**). However, BPF exposure resulted in FPDc shortening at higher concentrations (3.7% at 10 µM (q<0.05), 12.5% at 30 µM (q<0.0001), 32.4% at 100 µM (q<0.0001)). Treatment with BPS did not alter FPDc at any of the tested concentrations. FPDc restitution curves were generated by increasing the pacing frequency (1-2 Hz). A frequency-dependent effect was not observed for BPA in hiPSC-CM, although BPF exhibited a slight reverse-use dependency with FPDc shortening more prominent at slower frequencies (**Figure 4C, F**).

**Figure 4.**
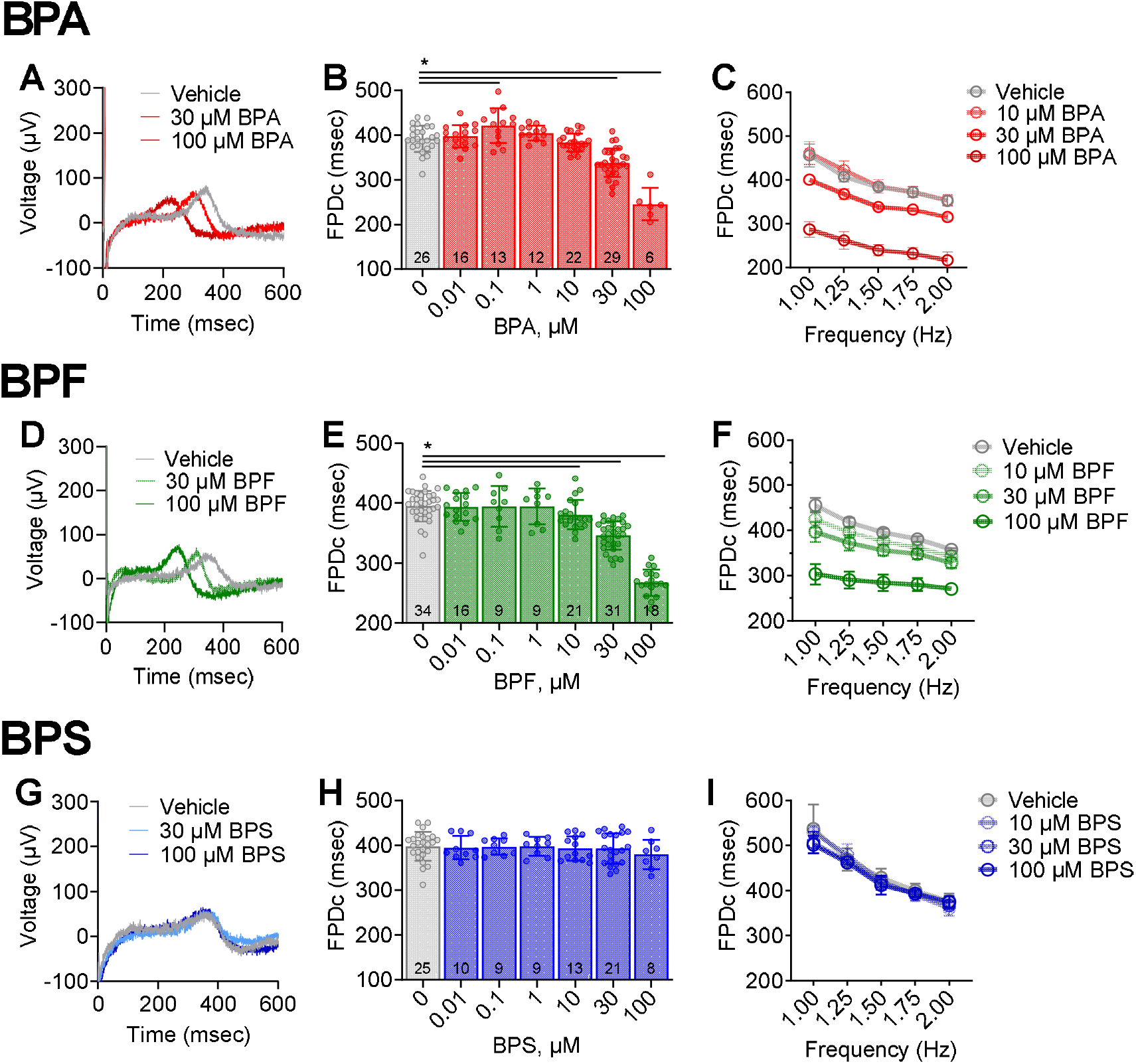
Cardiomyocyte field potential duration shortens with BPA or BPF exposure, but not BPS. **(A)** Representative traces of extracellular field potentials recorded from hiPSC-CM following acute exposure to vehicle, 30 μM, or 100 μM BPA. **(B)** Field potential duration (corrected using Frederica formula: ‘FPDc’) shortens with increasing BPA exposure; single pacing frequency (1.5 Hz). **(C)** FPDc restitution curve at multiple pacing frequencies (1-2 Hz). **(D)** Local field potential traces following exposure to vehicle, 30 μM, or 100 μM BPF. **(E)** FPDc shortens with increasing BPF exposure (1.5 Hz). **(F)** FPDc restitution curve (1-2 Hz). **(G)** Local field potential traces following exposure 30-100 μM BPS. **(H)** FPDc remains constant with increasing BPS exposure (1.5 Hz). **(I)** FPDc restitution curve (1-2 Hz). Values reported as mean ± SD. *q<0.05 as determined by ANOVA with multiple comparisons testing (0.1 FDR). Number of replicates indicated in each bar graph.

### BPA and BPF exposure slows heart rate

To aid in the translation of our *in vitro* findings, we quantified the acute effects of BPA, BPF, and BPS on cardiac electrophysiology using an *ex vivo* intact heart preparation. Heart preparations exhibited normal sinus rhythm when perfused with control KH media (327.7 ± 36.8 BPM) and KH media supplemented with vehicle (327.6 ± 40.3 BPM; **Figure 1C**). BPA exposure resulted in a measurable decline in heart rate, which may be partly attributed to calcium channel current inhibition. Sinus rate slowed by 16.6% (q<0.01) and 85.4% (q<0.0001) after exposure to 10 μM and 100 μM BPA, respectively (relative to baseline recording; **Figure 5A**,**B**). This depressive effect culminated in cessation of ventricular electrical activity in 62% of heart preparations treated with 100 μM BPA (**Figure 5D**). Heart rate slowing was immediate, yet reversible, as sinus rhythm recovered quickly after removal of BPA and replacement with control media perfusion. A non-monotonic BPA dose response relationship was not observed. Heart rate also slowed by 12.5% (q<0.001) after exposure to 100 μM BPF, the only dose to significantly affect automaticity (**Figure 5E**). Conversely, no significant change in sinus rhythm or rate were observed after exposure to BPS (**Figure 5G**).

**Figure 5.**
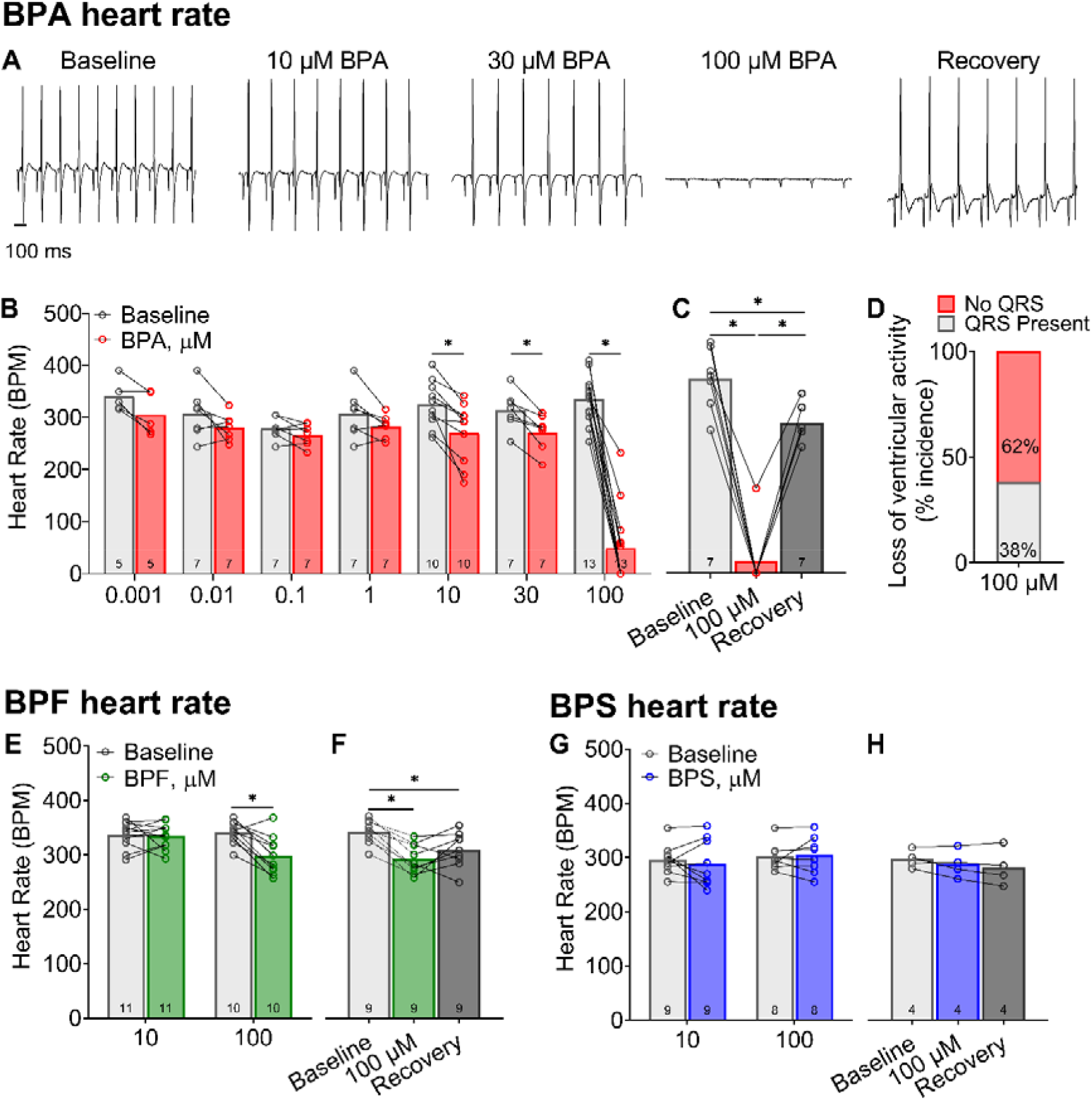
Heart rate slows in the presence of BPA or BPF exposure, but not BPS. **(A)** Representative ECG recordings from Langendorff-perfused hearts at baseline, acute (15 min) exposure to BPA, or recovery (100 μM BPA exposure, followed by 15 min washout). **(B)** BPA exposure results in sinus rate slowing, beginning at 10 μM BPA. **(C)** Heart rate slowing after 100 μM BPA exposure largely recovers after washout (15 min). **(D)** Heart rate measurements at high BPA doses were confounded by intermittent 3^rd^ degree heart block, with loss of ventricular electrical activity. **(E)** Heart rate slowing following BPF exposure occurred only at highest concentration tested (100 μM BPF). **(F)** Heart rate slowing after 100 μM BPF recovered slightly after washout. **(G**,**H)** BPS exposure had no discernable effect on heart rate. Values reported as mean ± SD. *q<0.05 as determined by RM-ANOVA with multiple comparisons testing (0.1 FDR). Number of replicates indicated in each bar graph.

### BPA and BPF exposure slows atrioventricular conduction

Heart preparations exhibited stable atrial and atrioventricular conduction with control media perfusion (15.3 ± 2.8 msec P duration, 33.6 ± 1.4 msec PR interval), and during perfusion with media supplemented with vehicle (16.3 ± 4.0 msec P duration, 35.3 ± 3.1 msec PR interval). Concurrent with heart rate slowing due to BPA exposure, we also observed significant lengthening of both the P duration and PR interval. A prolonged P duration was only observed at the highest BPA dose (39.8 ± 16.6 msec at 100 μM BPA, **Figure 6B**), whereas the PR interval progressively lengthened with higher BPA concentrations (**Figure 6A, D**). BPA exposure resulted in variable degrees of atrioventricular (AV) block, ranging from 1^st^ degree to intermittent 3^rd^ degree AV block (**Figure 6D**). Notably, the acute effect of BPA on AV conduction was reversible with a rapid recovery of the PR interval time after washout (**Figure 6E**). Atrial pacing was implemented, and AV conduction slowing persisted at multiple PCL (**Figure 6F**). BPF exposure also lengthened the PR interval, albeit the effect was much less pronounced (+19.6% 100 μM BPF). There was no observable change in AV conduction following BPS exposure (**Figure 6I**).

**Figure 6.**
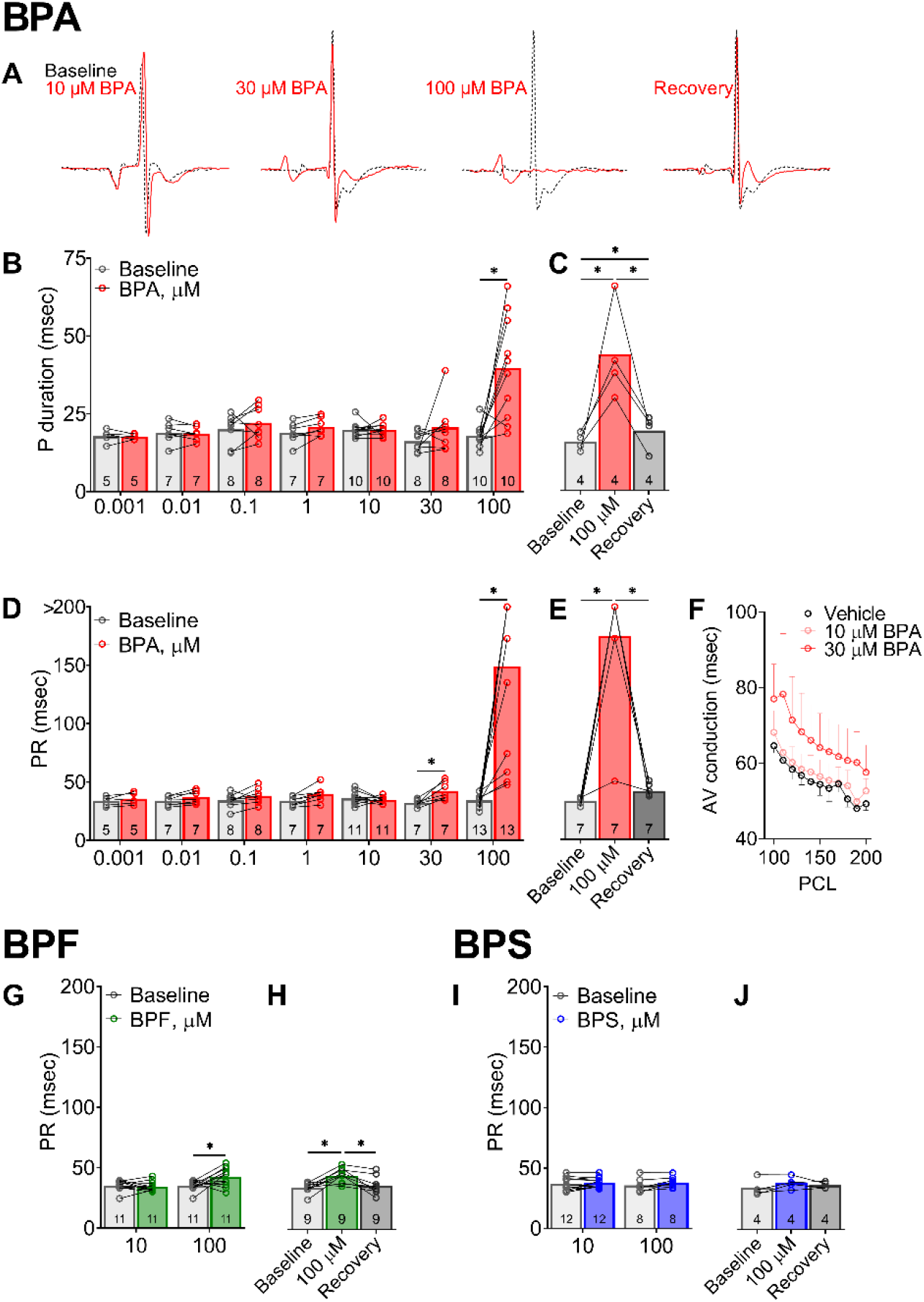
Atrial and atrioventricular conduction slows with BPA and BPF exposure, but not BPS. **(A)** Representative ECG waveform from Langendorff-perfused hearts at baseline, acute BPA exposure (15 min), or recovery (100 μM BPA exposure, followed by 15 min washout). Each waveform pair recorded from the same animal, before and after exposure. **(B)** P duration indicates longer atrial depolarization time at highest BPA dose (100 μM). **(C)** Slowed atrial conduction after 100 μM BPA exposure recovers after washout. **(D)** Progressive lengthening of PR duration indicates slowed AV conduction following 30-100 μM BPA exposure, often resulting in intermittent 3^rd^ degree heart block (denoted by data point >200 msec). **(E)** AV conduction slows after 100 μM BPA exposure and recovers after washout. **(F)** AV conduction slowing persists with external pacing to correct for heart rate. **(G)** Atrioventricular conduction slowing occurs only at highest BPF concentration (100 μM) and **(H)** recovers after washout. **(I**,**J)** BPS exposure had no discernable effect on atrioventricular conduction time. Values reported as mean ± SD. *q<0.05 as determined by RM-ANOVA with multiple comparisons testing (0.1 FDR). Number of replicates indicated in each bar graph. PCL= pacing cycle length

### BPA and BPF exposure increases atrioventricular nodal refractoriness

To further investigate slowed AV conduction in the presence of bisphenols, incremental atrial pacing was implemented to pinpoint the Wenckebach phenomenon. WBCL was comparable between control media perfusion (87 ± 6.2 msec) and during perfusion with vehicle (90 ± 6.5 msec). But, proved to be a highly sensitive parameter for bisphenol-induced slowing of AV conduction (**Figure 7A**,**B**). BPA exposure altered WBCL in a dose-dependent manner beginning at a low nanomolar concentration (+7.6% 0.01 μM BPA (q<0.1); +68% 100 μM BPA (q<0.0001)), suggesting a lengthening of the relative refractory period. Pinpointing an accurate WBCL after 100 μM BPA exposure was confounded by loss of capture at the slowest cycle length tested (150 msec; **Figure 7C**). BPF exposure had a moderate effect on WBCL, but only at the highest concentration tested (22.5% 100 μM BPF (q<0.0001)). BPS exposure did not alter WBCL, which was in agreement with PR interval measurements during sinus rhythm (**Figure 7G**). Extrastimulus pacing was also performed to measure the effective refractory period of the atrioventricular node. Similar to WBCL measurements, BPA exposure increased AV node refractoriness in a dose-dependent manner (**Figure 8A**,**B**), beginning with modest changes at a low nanomolar concentration (+9.2% 0.001 μM BPA (q<0.05)) and increasing thereafter (+95.7% 100 μM BPA (q<0.0001)). BPF exposure had a moderate effect on AVNERP, but only at the highest concentration tested (20.7% 100 μM BPF (q<0.0001)). BPS exposure did not alter AVNERP (**Figure 8G**).

**Figure 7.**
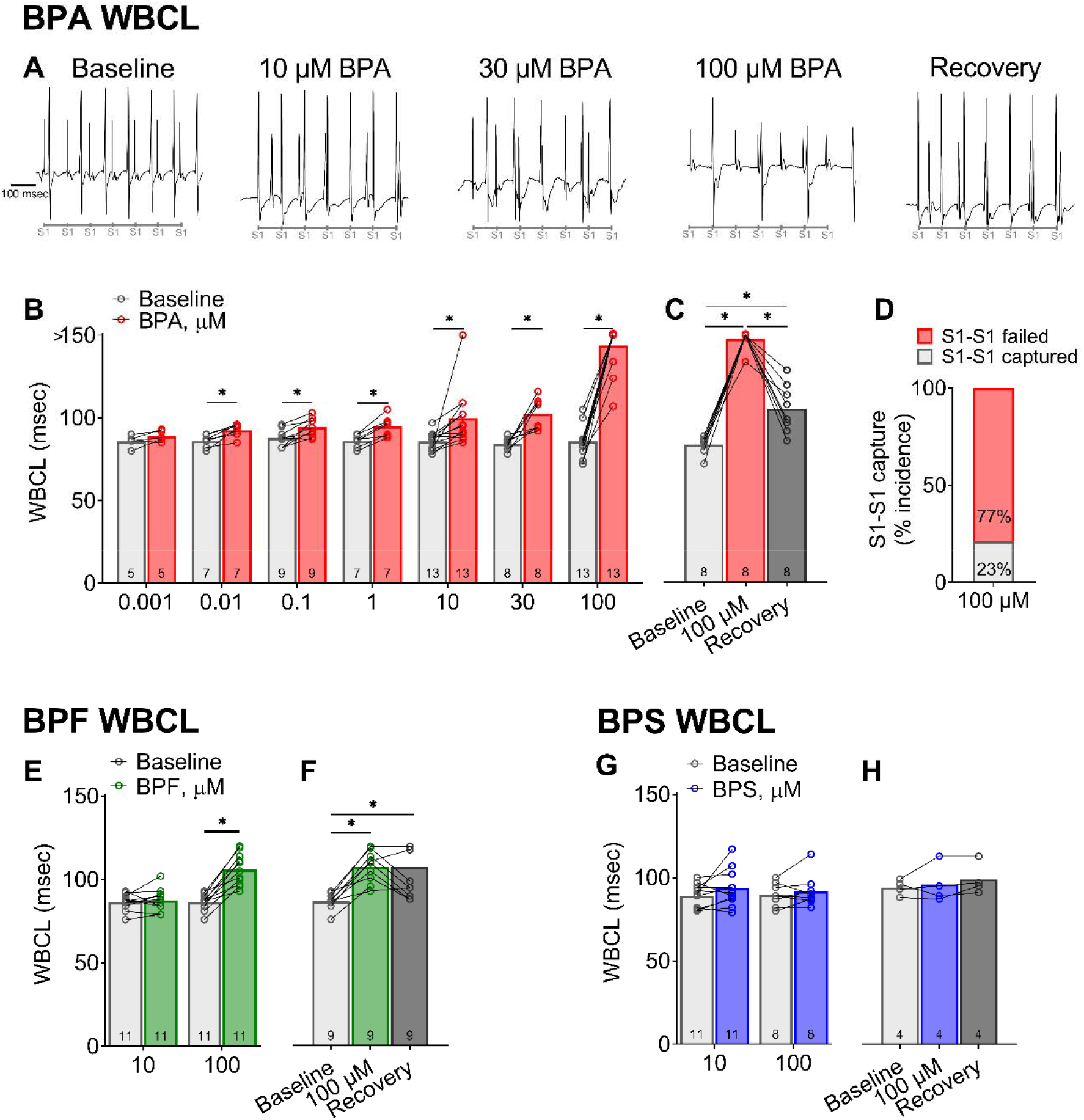
Atrial pacing highlights atrioventricular slowing after BPA or BPF, but not BPS exposure. **(A)** Representative ECG recordings during atrial pacing show failure to capture in BPA-treated hearts, indicating slowed atrioventricular conduction. Timing of S1-S1 pulses (90 msec) are indicated below. **(B)** Longer Wenckebach cycle length (WBCL) following exposure to BPA concentrations (0.01–100 μM), as compared with baseline. Note: Complete Heart block denoted by measurement >150 msec (longest S1 pacing interval tested). **(C)** Longer WBCL after 100 μM BPA exposure largely recovers after washout. **(D)** WBCL measurements at high BPA doses were confounded by complete heart block. **(E)** Only 100 μM BPF exposure results in longer WBCL. **(F)** Moderate recovery in atrioventricular conduction after BPF washout. **(G**,**H)** No change in WBCL was observed after exposure to BPS. Values reported as mean ± SD. *q<0.05 as determined by RM-ANOVA with multiple comparisons testing (0.1 FDR). Number of replicates indicated in each bar graph.

**Figure 8.**
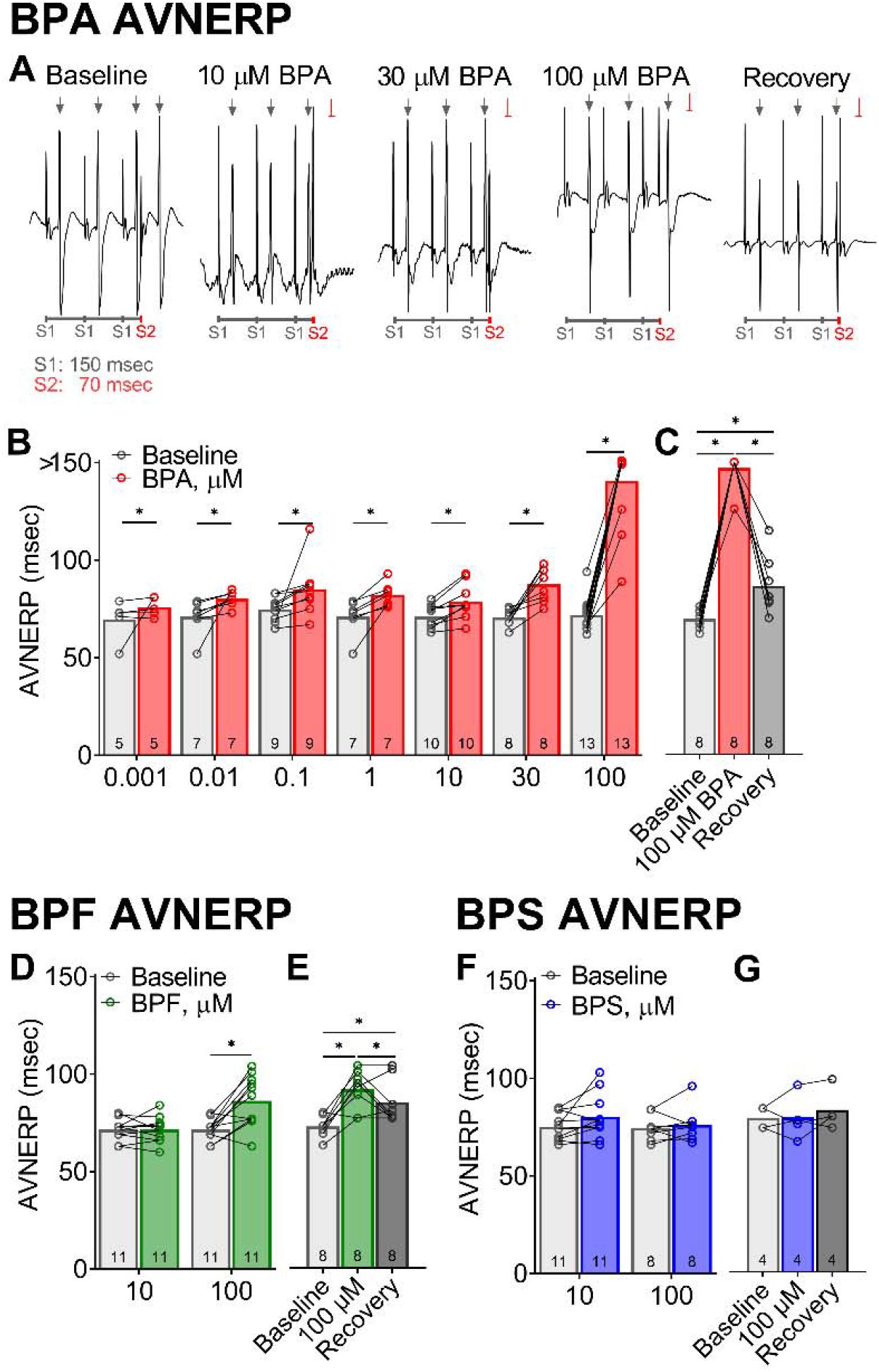
Increased atrioventricular nodal refractoriness after exposure to BPA and BPF, but not BPS. **(A)** Representative ECG recordings during atrial pacing show capture (_↓_) and failure to capture (-) in response to S1-S2 pacing (150, 70 msec). **(B)** Longer atrioventricular nodal effective refractory period (AVNERP) following exposure to BPA concentrations (1–100 μM), as compared with baseline. Note: Complete heart block denoted by measurement >150 msec (S1 pacing interval). **(C)** Longer AVNERP after 100 μM BPA exposure recovers after washout. **(D)** BPF exposure results in longer AVNERP, but only at highest dose tested. **(E)** Moderate recovery of AVNERP after BPF washout. **(F**,**G)** No change in AVNERP was observed after exposure to BPS. Values reported as mean ± SD. *q<0.05 as determined by RM-ANOVA with multiple comparisons testing (0.1 FDR). Number of replicates indicated in each bar graph.

## DISCUSSION

This is the first study to compare the direct effects of BPA, BPS, and BPF on cardiac electrophysiology using both *in vitro* and *ex vivo* cardiac preparations. In the described study, we demonstrate that BPA is the most potent inhibitor of sodium, calcium and potassium channel currents – as compared to the chemical alternatives, BPS and BPF. Using a range of concentrations that encompass environmental, clinical, and supraphysiological exposures, we found that BPA exerted the greatest effect on automaticity and atrioventricular conduction. Electrical disturbances were largely focused on nodal and atrioventricular conduction, with negligible effects on cardiac repolarization or arrhythmia susceptibility. Results of this study indicate that BPS is significantly less disruptive to cardiac electrophysiology and may serve as a safer chemical alternative for plastic medical products. It is important to note that in the context of industrial or clinical environments, individuals can present with urinary BPA concentrations that are exceedingly high – reaching 4-8 μM (Calafat et al. 2009; He et al. 2009; Wang et al. 2012). Notably, this study focused on acute bisphenol exposure and the direct impact on cardiac electrophysiology endpoints; as such, we did not investigate intracellular signaling or possible genomic effects of bisphenol chemicals. Additional mechanistic studies are required to fully elucidate the safety profile of bisphenol chemicals on cardiac electrical and mechanical function, and report on the chronic effects of bisphenol exposure.

### Bisphenol chemicals and calcium ion homeostasis

Of the bisphenol chemicals tested in this study, BPA was the most potent inhibitor of L-type calcium channels with an IC_50_ = 30.8 μM. This finding is in agreement with the literature, which reported an immediate, inhibitory effect of BPA on T-type calcium channels in HEK cells (IC_50_ = 6-33 μM, depending on channel subtype; (Michaela et al. 2014)). Similarly, Deutschmann, et al. reported that BPA rapidly and reversibly inhibited calcium current through L-, N-, P/Q-, R-, and T-type calcium channels in rat endocrine cells, dorsal root ganglion, cardiomyocytes, and transfected HEK cells (IC_50_ = 26-35 μM; (Deutschmann et al. 2013)). Studies suggest that the inhibitory effect of bisphenol chemicals on calcium channel current are influenced by the chemical structure and bridge between the two phenol rings, with reduced inhibitory effects anticipated for BPS and BPF (Deutschmann et al. 2013). In cardiac tissue, calcium channels play an important role in nodal cell depolarization, atrioventricular conduction, the plateau phase of the cardiac action potential, and contractility. Indeed, recent studies have shown that BPA can alter cardiac electrophysiology, likely through a calcium-dependent mechanism. Sinus bradycardia and delayed electrical conduction have been reported after BPA exposure, using *in vivo* and *ex vivo* models (Belcher et al. 2015; Patel et al. 2015; Posnack et al. 2014; Valokola et al. 2019). Patel et al. observed conduction slowing in BPA-exposed animals subjected to catecholamine stress, although this effect was limited to females (Patel et al. 2015). BPA-induced heart rate slowing was also reported in *in vivo* studies conducted by Belcher, et al., although the authors noted that this effect may be attributed to autonomic dysregulation (Belcher et al. 2015). In addition to electrophysiology disturbances, BPA-exposure has been shown to alter intracellular calcium handling, which can increase calcium leak from the sarcoplasmic reticulum (Yan et al. 2011), increase the incidence of calcium-mediated arrhythmias, and precipitate calcium amplitude alternans (Ramadan et al. 2018). Studies suggest that intracellular calcium handling may be influenced by posttranslational modifications of key calcium proteins, via an estrogen-mediated mechanism (Liang et al. 2014).

### Bisphenol chemicals and sodium channel current

Similar to calcium channel inhibition, we found that BPA was the most potent inhibitor of fast (I_Na-P_) and late (I_Na-L_) sodium channel (IC_50_ = 55.3 and 23.6 μM, respectively). This finding is in agreement with previously published studies, which reported that BPA blocks fast-voltage gated sodium channels in transfected HEK cells (IC_50_ = 25 μM; (O’Reilly et al. 2012)) and isolated dorsal root ganglion neurons (IC_50_ = 40 mM; (Wang et al. 2011)). These effects were rapid, reversible, and dose-dependent (Wang et al. 2011). Moreover, in isolated ganglion neurons, the described effects were attenuated with protein kinase A (PKA) or protein kinase C (PKC) inhibitors, suggesting an underlying protein-kinase dependent pathway. In cardiac tissue, the fast voltage-gated sodium channel (I_Na-P_; TTX-sensitive) is responsible for the action potential upstroke and blockade is likely to reduce the rate of depolarization and slow conduction velocity. Late-sodium channel current (TTX-insensitive) is active during the action potential plateau phase, and blockade is expected to shorten repolarization time, which in turn, reduces calcium channel current (Horváth et al. 2020). This highlights the importance of performing electrophysiology studies using cardiac models (e.g., human cardiomyocytes, isolated whole heart, in vivo studies) in conjunction with single channel studies, given the dynamic nature of cardiac electrophysiology.

### Bisphenol chemicals and estrogen-receptor signaling

Since BPA is classified as a xenoestrogen, alterations in cardiac function may also be attributed to its interaction with estrogen receptors. In the presented study, we utilized female cardiac preparations since previous reports have indicated that BPA-induced effects on calcium handling and electrical instabilities can be heightened in female animals, exacerbated in the presence of estradiol, and attenuated in ERβ knockout animals (Belcher et al. 2011; Yan et al. 2011). Further, studies have shown that 17β-estradiol alone can rapidly and reversibly inhibit sodium and calcium channels, in a concentration dependent manner (DY et al. 2002; Wang et al. 2013). Accordingly, the effects of BPA on cardiac electrophysiology may be mediated by direct interaction with ion channels at the cell membrane and/or intracellular signaling pathways precipitated by estrogen receptor binding. Of interest, BPS and BPF have also been shown to display estrogenic activity that is comparable to BPA (Kojima et al. 2019; Moreman et al. 2017). Although in the presented study, we identified clear differences in the potency of BPA, BPS, and BPF as it relates to ion channel inhibition and cardiac electrophysiology parameters.

### Cardiac safety profile of bisphenol A analogues

Biomonitoring studies have recently reported an uptick in BPS and BPF exposure in the general population, as manufacturers begin to phase out and replace BPA in some consumer and medical products. As an example, data from the 2013-2014 National Health and Nutrition Examination Survey (NHANES) detected BPA, BPS, and BPF in 96%, 90%, and 67% of urinary samples from the general population (Lehmler et al. 2018). Yet, very little is known about the effects of these substitute chemicals on cardiovascular health, and whether they offer a superior safety profile. Using a zebrafish model, Chen et al. reported that BPS exposure results in transcriptional changes that can increase inflammation, alter cardiac morphology, and *decrease* heart rate (Qiu et al. 2020). In a rodent cardiac model, BPS-treatment alone was shown to *increase* heart rate, while the addition of catecholamine stress increased the propensity for premature ventricular contractions and calcium-mediated triggered activity (Gao et al. 2015). The authors noted that the observed effect on cardiac electrophysiology was sex-specific and mediated via estrogen receptor-β signaling, which alters the phosphorylation status of key calcium handling proteins. Notably, the same group has reported nearly identical effects with BPA-treatment, which suggest that the two chemicals may act via a common mechanism (Yan et al. 2011). In a separate study by Ferguson, et al., BPS or BPA-treatment rapidly reduced mechanical function in heart preparations, but slightly different post-translational modifications were observed in myofilament proteins (Ferguson et al. 2019). Investigations into the cardiac effects of BPF are even more limited, with a single report noting a decrease in the heart rate of zebrafish following BPF-exposure (Mu et al. 2019). The current study was focused on cardiac electrophysiology outcomes, therefore, additional work is needed to assess the impact of BPA, BPF, and BPS on myocardial contractility.

To the best of our knowledge, our study is the first to compare the acute effects of BPA, BPS, and BPF exposure on cardiac electrophysiology. We aimed to identify the IC_50_ concentrations for BPA, BPS, and BPF on key cardiac ion channels highlighted by the CiPA initiative – and validate the effect of those concentrations on human cardiomyocyte and intact heart preparations. Collectively, we observed that BPA exposure has a more potent effect on cardiac electrophysiology, as compared to the chemical substitutes BPF and BPS. Our results suggest that BPS may be a safer chemical alternative, particularly for medical devices that are used to treat vulnerable patient populations that are at increased risk for bisphenol chemical exposure. Nevertheless, a few limitations to our study should be considered. Although we included a number of models in our study, further in-depth mechanistic work is necessary to fully elucidate the safety profile of bisphenol chemicals – including the impact on intracellular targets, genomic and proteomic expression profiles (sub-acute or chronic studies), and autonomic regulation (in vivo studies). There are also notable differences in cardiac electrophysiology between rodents and humans (e.g., ion channel expression, sinus rate, action potential morphology) which should be noted when considering the translation of experimental studies to humans.

## ACKNOWLEDGEMENTS

This work was supported by the National Institutes of Health (R01HL139472 to NGP), Children’s National Heart Institute, Sheikh Zayed Institute for Pediatric Surgical Innovation, and the Children’s National Research Institute. This publication was also supported by the Gloria and Steven Seelig family.

## Notes

**Conflict of interest**: None.

### Competing Interest Statement

The authors have declared no competing interest.

## REFERENCES

Asakura K, Hayashi S, Ojima A, Taniguchi T, Miyamoto N, Nakamori C, et al. Improvement of acquisition and analysis methods in multi-electrode array experiments with iPS cell-derived cardiomyocytes. J Pharmacol Toxicol Methods 75: 17–26.

Bae S, Hong Y-C. 2015. Exposure to bisphenol A from drinking canned beverages increases blood pressure: randomized crossover trial. Hypertension 65:313–319; doi:10.1161/HYPERTENSIONAHA.114.04261.

Bae S, Kim JH, Lim Y-H, Park HY, Hong Y-C. 2012. Associations of bisphenol A exposure with heart rate variability and blood pressure. Hypertension 60:786–793; doi:10.1161/HYPERTENSIONAHA.112.197715.

Bao W, Liu B, Rong S, Dai SY, Trasande L, Lehmler HJ. 2020. Association Between Bisphenol A Exposure and Risk of All-Cause and Cause-Specific Mortality in US Adults. JAMA Netw open 3:e2011620; doi:10.1001/jamanetworkopen.2020.11620.

Belcher SM, Chen Y, Yan S, Wang H-SS. 2011. Rapid Estrogen Receptor-Mediated Mechanisms Determine the Sexually Dimorphic Sensitivity of Ventricular Myocytes to 17beta-Estradiol and the Environmental Endocrine Disruptor Bisphenol A. Endocrinology 153:712–720; doi:10.1210/en.2011-1772.

Belcher SM, Gear RB, Kendig EL. 2015. Bisphenol A Alters Autonomic Tone and Extracellular Matrix Structure and Induces Sex-Specific Effects on Cardiovascular Function in Male and Female CD-1 Mice. Endocrinology 156:882–895; doi:10.1210/en.2014-1847.

Ben-Jonathan N, Steinmetz R. 1998. Xenoestrogens: The emerging story of bisphenol A. Trends Endocrinol Metab 9:124–128; doi:10.1016/S1043-2760(98)00029-0.

Birnbaum LS. 2012. Environmental chemicals: Evaluating low-dose effects. Environ Health Perspect 120; doi:10.1289/ehp.1205179.

Calafat AM, Kuklenyik Z, Reidy JA, Caudill SP, Ekong J, Needham LL. 2005. Urinary concentrations of bisphenol A and 4-Nonylphenol in a human reference population. Environ Health Perspect 113:391–395; doi:10.1289/ehp.7534.

Calafat AM, Weuve J, Ye X, Jia LT, Hu H, Ringer S, et al. 2009. Exposure to bisphenol A and other phenols in neonatal intensive care unit premature infants. Environ Health Perspect 117:639–644; doi:10.1289/ehp.0800265.

Chen D, Kannan K, Tan H, Zheng Z, Feng Y-L, Wu Y, et al. 2016a. Bisphenol Analogues Other Than BPA: Environmental Occurrence, Human Exposure, and Toxicity—A Review. Environ Sci Technol 50:5438–5453; doi:10.1021/acs.est.5b05387.

Chen IY, Matsa E, Wu JC. 2016b. Induced pluripotent stem cells: at the heart of cardiovascular precision medicine. Nat Publ Gr 13; doi:10.1038/nrcardio.2016.36.

Clements M. 2016. Multielectrode Array (MEA) Assay for Profiling Electrophysiological Drug Effects in Human Stem Cell-Derived Cardiomyocytes. In: Current Protocols in Toxicology. Vol. 68 of. John Wiley & Sons Inc.:, Hoboken, NJ, USA. 22.4.1-22.4.32.

Colatsky T, Fermini B, Gintant G, Pierson JB, Sager P, Sekino Y, et a. 2016. The Comprehensive in Vitro Proarrhythmia Assay (CiPA) initiative - Update on progress. J Pharmacol Toxicol Methods 81:15–20; doi:10.1016/j.vascn.2016.06.002.

Deutschmann A, Hans M, Meyer R, Häberlein H, Swandulla D. 2013. Bisphenol A inhibits voltage-activated Ca(2+) channels in vitro: mechanisms and structural requirements. Mol Pharmacol 83:501–511; doi:10.1124/mol.112.081372.

Duty SM, Mendonca K, Hauser R, Calafat AM, Ye X, Meeker JD, et al. 2013. Potential sources of bisphenol a in the neonatal intensive care unit. Pediatrics 131:483–489; doi:10.1542/peds.2012-1380.

Dy L, Yg C, Eb L, Kw K, Sy N, Th O, et al. 2002. 17Beta-estradiol inhibits high-voltage-activated calcium channel currents in rat sensory neurons via a non-genomic mechanism. Life Sci 70; doi:10.1016/S0024-3205(01)01534-X.

Edwards AG, Louch WE. 2017. Species-Dependent Mechanisms of Cardiac Arrhythmia: A Cellular Focus. Clin Med Insights Cardiol 11; doi:10.1177/1179546816686061.

Feiteiro J, Mariana M, Glória S, Cairrao E. 2018. Inhibition of L-type calcium channels by Bisphenol A in rat aorta smooth muscle. J Toxicol Sci 43:579–586; doi:10.2131/jts.43.579.

Ferguson M, Lorenzen-Schmidt I, Pyle WG. 2019. Bisphenol S rapidly depresses heart function through estrogen receptor-β and decreases phospholamban phosphorylation in a sex-dependent manner. Sci Rep 9:15948; doi:10.1038/s41598-019-52350-y.

Gao X, Liang Q, Chen Y, Wang H-SS. 2013. Molecular mechanisms underlying the rapid arrhythmogenic action of bisphenol A in female rat hearts. Endocrinology 154:4607–4617; doi:10.1210/en.2013-1737.

Gao X, Ma J, Chen Y, Wang H-S. 2015. Rapid responses and mechanism of action for low-dose bisphenol S on ex vivo rat hearts and isolated myocytes: evidence of female-specific proarrhythmic effects. Environ Health Perspect 123:571–8; doi:10.1289/ehp.1408679.

Gaynor JW, Ittenbach RF, Calafat AM, Burnham NB, Bradman A, Bellinger DC, et al. 2018. Perioperative Exposure to Suspect Neurotoxicants from Medical Devices in Newborns with Congenital Heart Defects. Ann Thorac Surg 107:567–572; doi:10.1016/j.athoracsur.2018.06.035.

Han C, Hong Y-C. 2016. Bisphenol A, Hypertension, and Cardiovascular Diseases: Epidemiological, Laboratory, and Clinical Trial Evidence. Curr Hypertens Rep 18:11; doi:10.1007/s11906-015-0617-2.

He Y, Miao M, Wu C, Yuan W, Gao E, Zhou Z, et al. 2009. Occupational exposure levels of bisphenol A among Chinese workers. J Occup Health 51:432–6; doi:10.1539/joh.o9006.

Hines CJ, Christianson AL, Jackson M V, Ye X, Pretty JR, Arnold JE, et al. 2018. An Evaluation of the Relationship among Urine, Air, and Hand Measures of Exposure to Bisphenol A (BPA) in US Manufacturing Workers. Ann Work Expo Heal 62:840–851; doi:10.1093/annweh/wxy042.

Horváth B, Hézső T, Kiss D, Kistamás K, Magyar J, Nánási PP, et al. 2020. Late Sodium Current Inhibitors as Potential Antiarrhythmic Agents. Front Pharmacol 11:413; doi:10.3389/fphar.2020.00413.

Huygh J, Clotman K, Malarvannan G, Covaci A, Schepens T, Verbrugghe W, et al. 2015. Considerable exposure to the endocrine disrupting chemicals phthalates and bisphenol-A in intensive care unit (ICU) patients. Environ Int 81:64–72; doi:10.1016/j.envint.2015.04.008.

Iribarne-Durán LM, Artacho-Cordón F, Peña-Caballero M, Molina-Molina JM, Jiménez-Díaz I, Vela-Soria F, et al. 2019. Presence of bisphenol a and parabens in a neonatal intensive care unit: An exploratory study of potential sources of exposure. Environ Health Perspect 127; doi:10.1289/EHP5564.

Jaimes R, McCullough D, Siegel B, Swift L, McInerney D, Hiebert J, et al. 2019. Plasticizer Interaction with the Heart: Chemicals Used in Plastic Medical Devices Can Interfere with Cardiac Electrophysiology. Circ Arrhythmia Electrophysiol 12; doi:10.1161/CIRCEP.119.007294.

Koch HM, Calafat AM. 2009. Human body burdens of chemicals used in plastic manufacture. Philos Trans R Soc B Biol Sci 364:2063–2078; doi:10.1098/rstb.2008.0208.

Kojima H, Takeuchi S, Sanoh S, Okuda K, Kitamura S, Uramaru N, et al. 2019. Profiling of bisphenol A and eight of its analogues on transcriptional activity via human nuclear receptors. Toxicology 413:48–55; doi:10.1016/J.TOX.2018.12.001.

Lehmler HJ, Liu B, Gadogbe M, Bao W. 2018. Exposure to Bisphenol A, Bisphenol F, and Bisphenol S in U.S. Adults and Children: The National Health and Nutrition Examination Survey 2013-2014. ACS Omega 3:6523–6532; doi:10.1021/acsomega.8b00824.

Liang Q, Gao X, Chen Y, Hong K, Wang H-SS. 2014. Cellular mechanism of the nonmonotonic dose response of bisphenol A in rat cardiac myocytes. Environ Health Perspect 122:601– 608; doi:10.1289/ehp.1307491.

Mantegazza M, Franceschetti S, Avanzini G. 1998. Anemone toxin (ATX II)-induced increase in persistent sodium current: effects on the firing properties of rat neocortical pyramidal neurones. J Physiol 507 (Pt 1): 105–16.

Melzer D, Osborne NJ, Henley WE, Cipelli R, Young A, Money C, et al. 2012. Urinary bisphenol A concentration and risk of future coronary artery disease in apparently healthy men and women. Circulation 125:1482–1490; doi:10.1161/CIRCULATIONAHA.111.069153.

Melzer D, Rice NE, Lewis C, Henley WE, Galloway TS. 2010. Association of urinary bisphenol a concentration with heart disease: evidence from NHANES 2003/06. PLoS One 5:e8673– e8673; doi:10.1371/journal.pone.0008673.

Michaela P, Mária K, Silvia H, Ľubica L, L’ubica L. 2014. Bisphenol A Differently Inhibits CaV3.1, Ca V3.2 and Ca V3.3 Calcium Channels. Naunyn Schmiedebergs Arch Pharmacol 387; doi:10.1007/s00210-013-0932-6.

Moon MK. 2019. Concern about the Safety of Bisphenol A Substitutes. Diabetes Metab J 43:46–48; doi:10.4093/dmj.2019.0027.

Moreman J, Lee O, Trznadel M, David A, Kudoh T, Tyler CR. 2017. Acute Toxicity, Teratogenic, and Estrogenic Effects of Bisphenol A and Its Alternative Replacements Bisphenol S, Bisphenol F, and Bisphenol AF in Zebrafish Embryo-Larvae. Environ Sci Technol 51:12796–12805; doi:10.1021/ACS.EST.7B03283.

Mu X, Liu J, Yuan L, Yang K, Huang Y, Wang C, et al. 2019. The mechanisms underlying the developmental effects of bisphenol F on zebrafish. Sci Total Environ 687; doi:10.1016/J.SCITOTENV.2019.05.489.

O’Reilly AO, Eberhardt E, Weidner C, Alzheimer C, Wallace BA, Lampert A. 2012. Bisphenol a binds to the local anesthetic receptor site to block the human cardiac sodium channel. PLoS One 7:e41667–e41667; doi:10.1371/journal.pone.0041667.

Patel BB, Raad M, Sebag IA, Chalifour LE. 2015. Sex-specific cardiovascular responses to control or high fat diet feeding in C57bl/6 mice chronically exposed to bisphenol A. Toxicol Reports 2:1310–1318; doi:10.1016/j.toxrep.2015.09.008.

Posnack NG. 2014. The Adverse Cardiac Effects of Di(2-ethylhexyl)phthalate and Bisphenol A. Cardiovasc Toxicol 14:339–357; doi:10.1007/s12012-014-9258-y.

Posnack NG, Brooks D, Chandra A, Jaimes R, Sarvazyan N, Kay MW. 2015. Physiological response of cardiac tissue to Bisphenol A: alterations in ventricular pressure and contractility. Am J Physiol Heart Circ Physiol 309:H267–H275; doi:10.1152/ajpheart.00272.2015.

Posnack NG, Jaimes R, Asfour H, Swift LM, Wengrowski AM, Sarvazyan N, et al. 2014. Bisphenol A Exposure and Cardiac Electrical Conduction in Excised Rat Hearts. Environ Health Perspect 122:384–90; doi:10.1289/ehp.1206157.

PR newswire. 2016. Global Bisphenol-A Market Overview 2016-2022. Available: https://www.prnewswire.com/news-releases/global-bisphenol-a-market-overview-2016-2022---market-is-projected-to-reach-us225-billion-by-2022-up-from-156-billion-in-2016---research-and-markets-300303934.html [accessed 31 January 2021].

Qiu W, Chen B, Greer JB, Magnuson JT, Xiong Y, Zhong H, et al. 2020. Transcriptomic Responses of Bisphenol S Predict Involvement of Immune Function in the Cardiotoxicity of Early Life-Stage Zebrafish (Danio rerio). Environ Sci Technol 54:2869–2877; doi:10.1021/acs.est.9b06213.

Ramadan M, Cooper B, Posnack NG. 2020. Bisphenols and phthalates: Plastic chemical exposures can contribute to adverse cardiovascular health outcomes. Birth Defects Res; doi:10.1002/bdr2.1752.

Ramadan M, Sherman M, Jaimes R, Chaluvadi A, Swift L, Posnack NGNG, et al. 2018. Disruption of neonatal cardiomyocyte physiology following exposure to bisphenol-a. Sci Rep 8:7356; doi:10.1038/s41598-018-25719-8.

Ribeiro E, Ladeira C, Viegas S. 2017. Occupational Exposure to Bisphenol A (BPA): A Reality That Still Needs to Be Unveiled. Toxics 5; doi:10.3390/toxics5030022.

Sager PT, Gintant G, Turner JR, Pettit S, Stockbridge N. 2014. Rechanneling the cardiac proarrhythmia safety paradigm: A meeting report from the Cardiac Safety Research Consortium. Am Heart J 167:292–300; doi:10.1016/j.ahj.2013.11.004.

Shelby MD. 2008. NTP-CERHR monograph on the potential human reproductive and developmental effects of bisphenol A. NTP CERHR MON v, vii–ix, 1-64 passim.

Talajic M, Nattel S. 1986. Frequency-dependent effects of calcium antagonists on atrioventricular conduction and refractoriness: Demonstration and characterization in anesthetized dogs. Circulation 74:1156–1167; doi:10.1161/01.CIR.74.5.1156.

Testai E, Hartemann P, Rodríguez-Farre E, Rastogi SC, Bustos J, Gundert-Remy U, et al. 2016. The safety of the use of bisphenol A in medical devices. Regul Toxicol Pharmacol 79:106–107; doi:10.1016/j.yrtph.2016.01.014.

Trasande L. 2017. Exploring regrettable substitution: replacements for bisphenol A. Lancet Planet Heal 1:e88–e89; doi:10.1016/S2542-5196(17)30046-3.

Valokola MG, Karimi G, Razavi BM, Kianfar M, Jafarian AH, Jaafari MR, et al. 2019. The protective activity of nanomicelle curcumin in bisphenol A-induced cardiotoxicity following subacute exposure in rats. Environ Toxicol 34:319–329; doi:10.1002/tox.22687.

Vandenberg LN. 2014. Non-monotonic dose responses in studies of endocrine disrupting chemicals: Bisphenol a as a case study. Dose-Response 12:259–276; doi:10.2203/dose-response.13-020.Vandenberg.

Vandenberg LN, Chahoud I, Heindel JJ, Padmanabhan V, Paumgartten FJR, Schoenfelder G. 2010. Urinary, circulating, and tissue biomonitoring studies indicate widespread exposure to bisphenol A. Environ Health Perspect 118:1055–70; doi:10.1289/ehp.0901716.

Vandenberg LN, Hauser R, Marcus M, Olea N, Welshons W V. 2007. Human exposure to bisphenol A (BPA). Reprod Toxicol 24:139–177; doi:10.1016/j.reprotox.2007.07.010.

Wang F, Hua J, Chen M, Xia Y, Zhang Q, Zhao R, et al. 2012. High urinary bisphenol A concentrations in workers and possible laboratory abnormalities. Occup Environ Med 69:679–684; doi:10.1136/oemed-2011-100529.

Wang Q, Cao J, Hu F, Lu R, Wang J, Ding H, et al. 2013. Effects of estradiol on voltage-gated sodium channels in mouse dorsal root ganglion neurons. Brain Res 1512:1–8; doi:10.1016/j.brainres.2013.02.047.

Wang Q, Cao J, Zhu Q, Luan C, Chen X, Yi X, et al. 2011. Inhibition of voltage-gated sodium channels by bisphenol A in mouse dorsal root ganglion neurons. Brain Res 1378:1–8; doi:10.1016/j.brainres.2011.01.022.

Yan S, Chen Y, Dong M, Song W, Belcher SM, Wang HS. 2011. Bisphenol A and 17β-estradiol promote arrhythmia in the female heart via alteration of calcium handling. PLoS One 6:e25455; doi:10.1371/journal.pone.0025455.

